# A novel ribosomal protein S20 variant in a family with unexplained colorectal cancer and polyposis

**DOI:** 10.1101/2019.12.16.877084

**Authors:** Bryony A. Thompson, Angela K. Snow, Cathryn Koptiuch, Wendy K. Kohlmann, Ryan Mooney, Sara Johnson, Chad Huff, Yao Yu, Craig C. Teerlink, Bing-Jian Feng, Deborah W. Neklason, Lisa A. Cannon-Albright, Sean V. Tavtigian

## Abstract

Colorectal cancer (CRC) has a large hereditary component, which is only partially explained by known genetic causes. Recently, variants in ribosomal protein S20 (*RPS20*, [OMIM: 603682]) were identified in a family with familial CRC type X and in a CRC cancer case-control screen. This study describes a novel splice donor variant in *RPS20*, NM_001023.3:c.177+1G>A. It segregates with CRC [OMIM: 114500] and polyposis [HP: 0200063] within the proband’s family. Reverse transcription-polymerase chain reaction (RT-PCR) confirms the variant results in two aberrantly-spliced transcripts that are absent in controls. The location of the novel *RPS20* variant is near two previously-reported truncating *RPS20* variants associated with CRC. DNA from colon adenocarcinoma showed no evidence of loss-of-heterozygosity, supporting a haploinsufficiency or dominant negative disease mechanism. These findings support designation of *RPS20* as a CRC predisposition gene, and expand the phenotypic spectrum of *RPS20* truncating variants to include polyposis.

## INTRODUCTION

With an estimated 132,700 new diagnoses and 49,700 deaths in 2015, colorectal cancer (CRC) remains one of the most common cancers in the US^1^. Family and twin studies estimate 20 - 30% of CRC is due to heritable factors^2-4^. Great strides have been made in understanding genetic factors that underlie CRC risk^5^. However, known genetic factors only explain about one third of familial risk of CRC^3^.

Ribosomal protein S20 (*RPS20*, [OMIM: 603682]) is a highly-conserved component of the 40S small ribosomal subunit that also functions in 18S rRNA processing^6^. Two studies have reported a link between truncating germline variants in *RPS20* and CRC. One frameshift variant (c.147dupA, p.Val50SerfsTer23) was reported to segregate with CRC in a large Finnish family^7^. A second frameshift variant (c.181_182delTT, p.Leu61GlufsTer11) as well as a predicted deleterious missense variant (c.160G>C, p.Val54Leu) were identified in a single CRC case each during whole exome sequencing of 863 CRC cases; no rare missense or truncating variants were identified in 1604 controls^8^.

Here we report a novel germline splice donor variant in *RPS20* (c.177+1G>A) identified in a CRC case from a case-control screen. This variant segregates with CRC and polyposis in additional family members.

## METHODS

### Editorial Policies and Ethical Considerations

Research was approved by the University of Utah Institutional Review Board. Individuals provided informed consent for genetic research under University of Utah protocols prior to collection of DNA, RNA, and/or FFPE tissue samples as well as medical record information.

### Case-Control Sequencing

Detailed methods for the case-control screen are provided in the Supplemental Methods. Briefly, DNA libraries were prepared from 746 CRC cases and 1,525 cancer-free controls using the Ovation Ultralow Library System v2 (NUGEN # 0347-A01). Library enrichment with a custom 196 gene panel (NimbleGen #06-471-722-001) was performed using the SeqCap EZ Reagent Kit Plus v2 (NimbleGen #06-953-247-001). Captured libraries were sequenced on an Illumina HiSeq2000 using the HiSeq 101 Cycle Paired-End protocol. Variant discovery was conducted using the Broad Institute’s Genome Analysis Tool Kit Best Practices germline workflows^9^.

### Statistics and Bioinformatics

Gene-disease association tests in the case-control cohort were performed using PERCH ^10^ and VAAST^11^. A significance level of P = 2.5 × 10^−4^ was set based on Bonferroni correction of P = 0.05^12^. MaxEntScan was used to predict alterations in splicing for *RPS20*^13^. RVsharing software was used to calculate the probability of the observed transmission of *RPS20* c.177+1G>A in the pedigree^14^.

### *RPS20* Genotyping

*RPS20* variant positions correspond to RefSeq NM_001023.3. DNA was extracted from whole blood (proband) using Qiagen AutoPure LS (Qiagen #2016391) or from saliva (additional pedigree members) using Oragene•DISCOVER (DNA Genotek #OGR-500). The variant-containing region was PCR amplified and Sanger sequenced (Supplemental Table S1).

### *RPS20* Reverse Transcription-PCR (RT-PCR)

Total RNA for the proband was extracted from whole blood using the PAXgene Blood RNA system (Qiagen #762164, #762165). Total RNA for the two controls was extracted from whole blood using a Qiagen miRNAeasy Mini Kit (Qiagen #217004). cDNA was produced using SuperScript IV VILO (Invitrogen #11756050), then PCR amplified and analyzed by gel electrophoresis (Supplemental Table S1). Three bands present for the proband (370bp, 444bp, 591bp) were cut from the agarose gel and incubated at 4°C overnight in 10mM Tris, 1mM EDTA, pH 8.0. A 5 µL aliquot of the resulting solution was used as a template in a second PCR reaction. These PCR products were treated with ExoSAP-IT (Affymetrix #78200), then analyzed by Sanger sequencing (Supplemental Table S1).

### Loss-of-Heterozygosity Analysis

DNA from individual II-3 (Figure 1) was extracted from normal colon and adenocarcinoma FFPE tissue. The *RPS20* variant was PCR amplified (Supplemental Table S1), and PCR bands were extracted using a QIAquick Gel Extraction kit (Qiagen #28704). DNA was treated with ExoSAP-IT prior to Sanger sequencing. Chromatographs from normal and tumor samples were compared by visual inspection.

**Figure 1.**
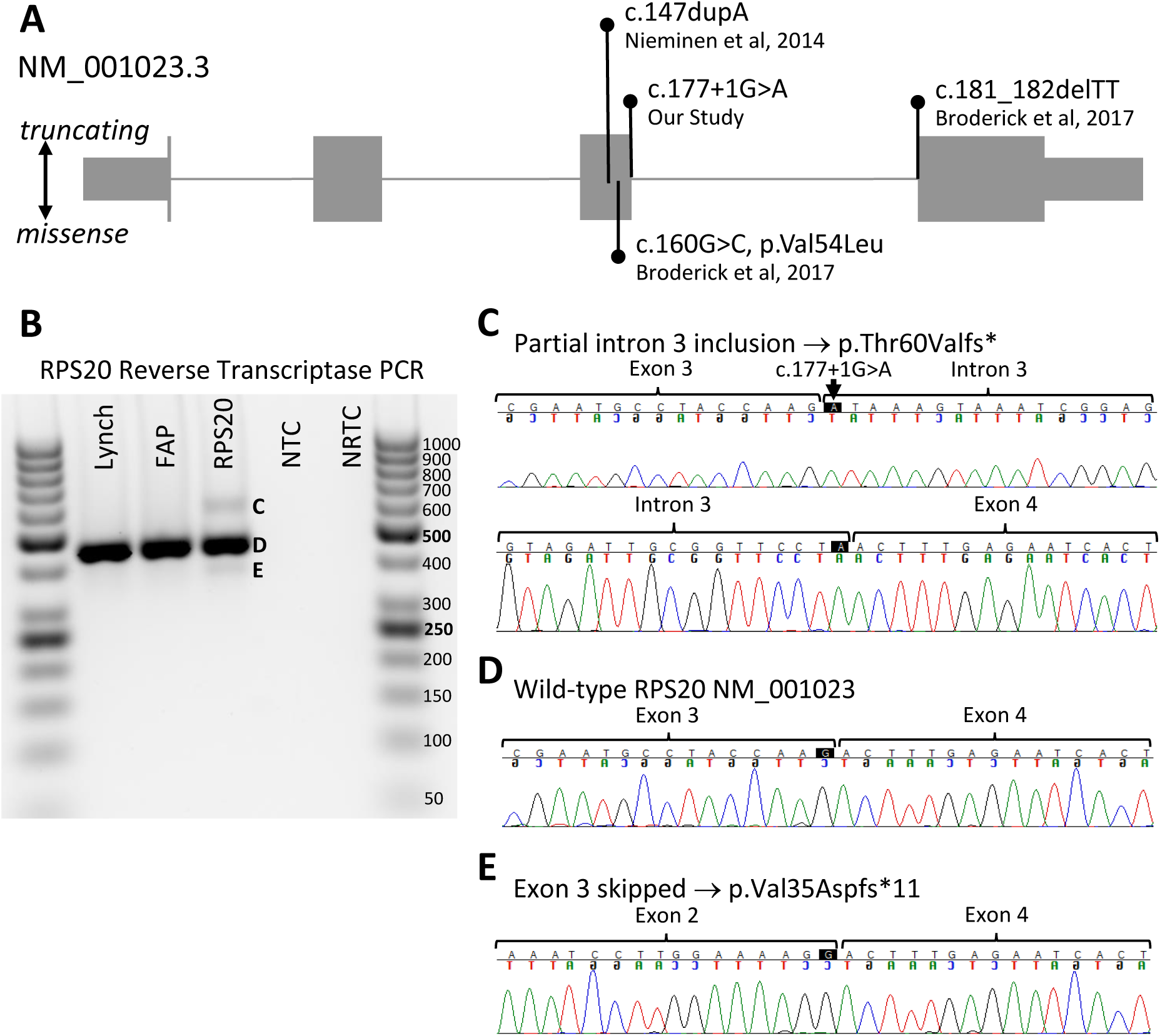
Confirmation of splicing defect. A) *RPS20* NM_001023.3 transcript with variants linked to CRC. c.147dupA (Nieminen et al, 2014). c.177+1G>A (this study). c.181_182delTT (Broderick et al, 2017). B) RT-PCR of *RPS20* from Lynch syndrome, FAP, and *RPS20* c.177+1G>A subjects. FAP (familial adenomatous polyposis); NTC (no template control); NRTC (no reverse transcriptase control). C) 591bp band corresponding to partial intron 3 inclusion. Black arrow indicates c.177+1G>A variant position. Note the chromatogram shows only adenine. D) 444bp band corresponding to wild-type NM_001023.3 sequence. E) 370bp band corresponding to exon 3 skipping.

## RESULTS

Germline DNA from CRC cases and controls was sequenced for 59 published (including *RPS20*) and 137 candidate cancer susceptibility genes (Supplemental Table S2). None of the 196 genes have a significant association with CRC after correction for multiple testing. However, a single rare variant in the canonical splice donor dinucleotide of *RPS20* intron 3 was identified in 1/746 CRC case and 0/1525 controls. Notably, the two previously reported *RPS20* variants are also truncating variants located near the junction of exons 3 and 4 (Figure 1A). Predicted loss-of-function variants in *RPS20* are rare, with only twelve carriers observed in the gnomAD non-cancer database (Supplemental Table S3)^15^. The G>A transition in the conserved splice donor site is predicted to affect mRNA splicing. RT-PCR was targeted to c.177+1G>A for the proband and for two controls with CRC syndromes (Lynch syndrome and familial adenomatous polyposis (FAP)) unrelated to *RPS20*. RT-PCR revealed three splice products in the proband’s mRNA and a single splice product in the two controls (Figure 1B-E). The splice products correspond to wild-type *RPS20* (Figure 1D), exon 3 skipping (r.104_177; p.Val35AspfsTer13, Figure 1E), and partial inclusion of intron 3 due to activation of a cryptic donor (r.177_178ins177+1_177+147; p.Thr60ValfsTer3, Figure 1C). The sequence of the intron 3 cryptic splice donor site, CTAgtaagt, is similar to the canonical human splice donor sequence (C/A)AGgt(a/g)agt^16^. It is predicted to be slightly weaker than the wild-type donor site for exon 3 (MaxEntScan scores - WT donor: 9.06; cryptic donor: 8.78), and is exclusively detected in the proband (Figure 1B).

The proband’s pedigree shows CRC in three of four siblings, the earliest at age 38 (Figure 2). Sanger sequencing revealed the c.177+1G>A variant’s presence in two siblings with CRC (II-1 and II-3) and one sibling with polyposis (II-4; Figure 2 and Table 1). The proband’s mother (I-2) is negative for the variant, suggesting it was inherited from the father (I-1), who had CRC at age 58.

**Table 1:** Clinical and Genetic Findings of Pedigree.

| ID | Sex | RPS20 status | Polyps at colonoscopy (age, source) | Cancer diagnosis (age, source) |
| --- | --- | --- | --- | --- |
| I-1 | M | ND |  | Colon (58, FR) |
| I-2 | F | WT | 3- AD 5mm (72, MR)<br>3- HP (72, MR)<br>13- HP 2-4mm (75, MR)<br>1- AD 3mm (80, MR)<br>4- HP 1mm (80, MR) |  |
| II-1 | F | c.177+1G>A | 0- (44, MR)<br>1- HP 1-2mm (49, MR)<br>1- TA 4-5mm (49, MR)<br>1- HP 5mm (50, MR)<br>0- (51, MR)<br>0- (53, MR)<br>1- TA 3-4mm (54, MR)<br>1- SP 6mm (54, MR)<br>1- TA <6mm (56, MR)<br>2- TA <6mm (57, MR)<br>1- U <6mm (57, MR)<br>1- SP 8mm (58, MR)<br>1- U <6mm (58, MR)<br>1- SP 3mm (59, MR)<br>1- U<6mm (60, MR)<br>3- TA (61, MR)<br>0- (62, MR) | Carcinoid – transverse (57, MR)<br>Colon (61, MR) |
| II-2 | M | c.177+1G>A | 0- (42, MR)<br>1- SP 3mm (44, MR)<br>0- (46, MR)<br>1- HP 3mm (47 MR)<br>2- HP 2-3mm (48, MR)<br>1- TA 6mm (50 MR)<br>2- U 2mm (50, MR)<br>1- HP 5mm (50, MR) | Colon – transverse (38, MR)<br>Colon recurrence (41, MR) |
| II-3 | M | c.177+1G>A | 0- (48, MR)<br>1- TA 5mm (57, MR)<br>1- TA 2mm (57, MR)<br>0- (58, MR) | Colon – splenic flexure (56, MR)<br>Colon recurrence – anastomotic site (58, MR) |
| II-4 | M | c.177+1G>A | 3- TA (53, MR)<br>1- TA 3mm (54, MR)<br>11- TA (57, MR)<br>0- (58, MR)<br>1- HP (59, MR)<br>0- (60, MR)<br>1- TA 7mm (61, MR)<br>1- TA 3mm (62, MR)<br>0- (63, MR) |  |
| III-1 | F | c.177+1G>A | 2- TA 2 mm (42, MR)<br>1- TA 2 mm (45, MR) |  |
| III-2 | M | c.177+1G>A | ND |  |
| III-3 | M | c.177+1G>A | No colonoscopies |  |
| III-4 | M | WT | 0- (24, SR) |  |
| III-5 | F | c.177+1G>A | No colonoscopies |  |

M (male); F (female); WT (wild type); ND (no data available); MR (confirmed by medical record); SR (self-reported); FR (family-reported); TA (tubular adenoma); AD (unspecified adenoma); HP (hyperplastic polyp); SP (sessile polyp); U (unknown polyp type). RPS20 reference transcript NM\_001023.3

**Figure 2.**
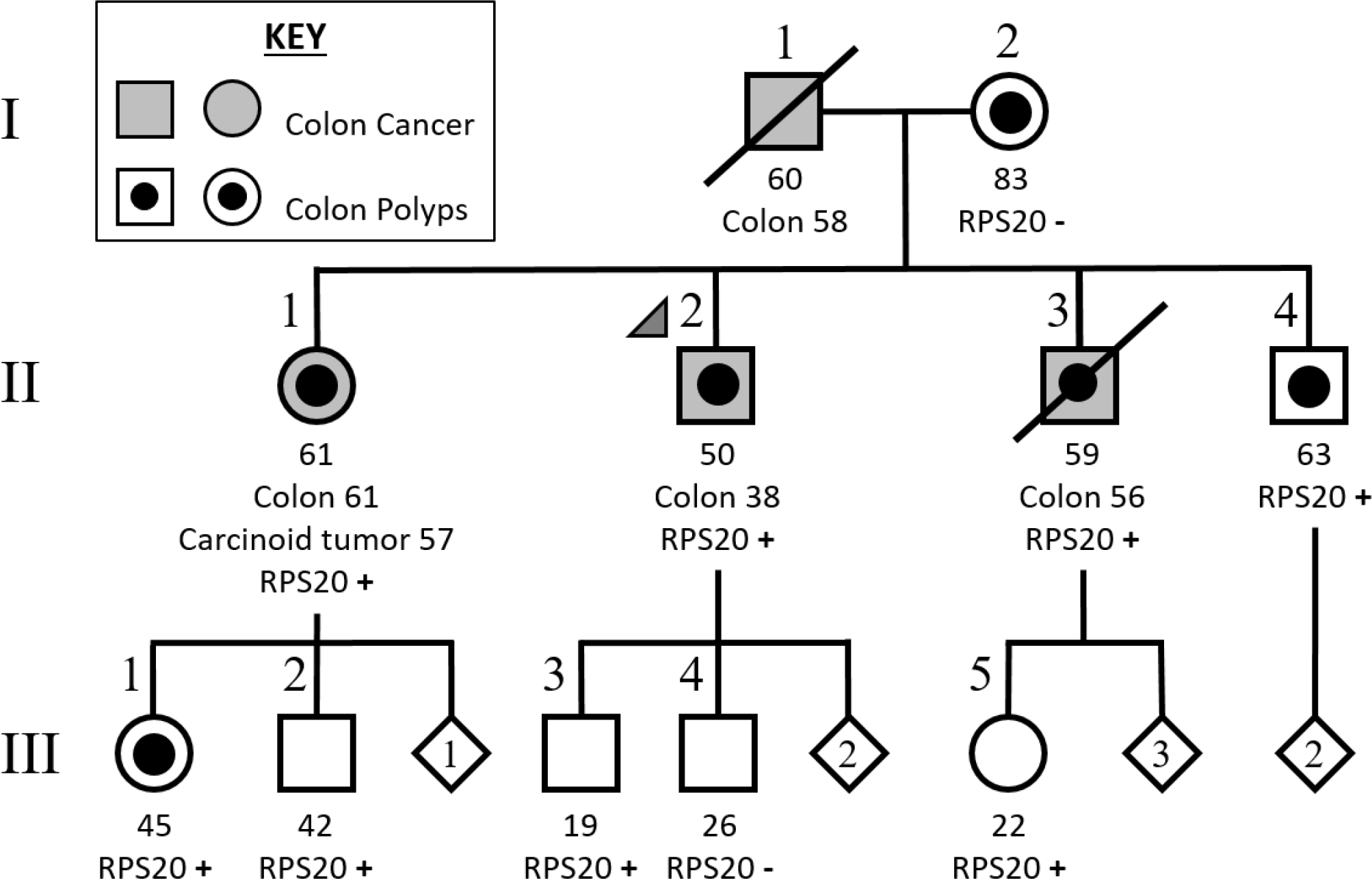
Pedigree of *RPS20* NM_001023.3:c.177+1G>A family. Arrow indicates proband. Age of individual at time of study or death indicated below square/circle. Cancers are reported by type and age at diagnosis. RPS20- indicates wild type sequence. RPS20+ indicates c.177+1G>A variant. Diamonds include number of additional siblings.

CRC tumor and normal tissue was available for individual II-3. There is no evidence of loss-of-heterozygosity comparing tumor and germline DNA, consistent with a previous report on tumors from *RPS20* c.147dupA carriers^7^ (Supplemental Figure S1).

## DISCUSSION

This work provides supporting evidence for an association between truncating *RPS20* variants and CRC. Mean age of first CRC diagnosis in this family is 53 years (50 years excluding I-1 whose *RPS20* status is unconfirmed), which is similar to the mean age of 52.3 years reported in the pedigree from Finland^7^. Unlike the Finnish *RPS20* family, which was ascertained as having hereditary nonpolyposis colorectal cancer^7^, two carriers in our family had ≥10 polyps (Table 1). A variable polyp phenotype, including individuals with no polyps, is observed in polyposis syndromes such as attenuated FAP (AFAP) and polymerase proofreading associated polyposis (PPAP) ^17,18^. Identification of additional affected families is required to better define the polyposis and cancer phenotype associated with pathogenic *RPS20* variants.

Nieminen et al. proposed haploinsufficiency as the likely mechanism of carcinogenesis for *RPS20*^7^. However, a dominant negative mechanism was not ruled out. Nonsense mediated decay (NMD) is not predicted to occur if the premature termination codon (PTC) occurs in the last exon or within the last 50 bp of the penultimate exon^19,20^. Notably, all reported truncating *RPS20* variants linked to CRC are near the exon 3 / 4 boundary, and are predicted to introduce PTCs within the last exon. If translated, all the respective proteins would contain an intact exon 2. Three of the proteins would have the same 10 amino acid C-terminal sequence (Supplemental Figure S2), with the final four amino acids conforming to the Class I PDZ ligand consensus sequence X[T/S]XΦ, where Φ is a hydrophobic residue and X is any amino acid ^21^. Additional families would help elucidate the association between variant type/location and CRC predisposition.

This study is limited by small family size and the young age of the lower generation compared to the median age-of-onset of CRC. Because of this, the association between *RPS20* c.177+1G>A and CRC does not reach statistical significance in the family (RVsharing pvalue 0.1429). The case-control analysis of *RPS20* also does not show a significant association (PERCH pvalue 0.076; VAAST pvalue 0.0461). Despite these limitations, given the rarity of predicted loss-of-function *RPS20* variants in the population^15^, it is important that our results are evaluated in conjunction with existing and emerging data. Future studies should pay particular attention to PTCs that would escape NMD, encode proteins with exons 1-3 mostly intact, and perhaps include a +1 frameshift peptide from exon 4.

## ACKNOWLEDGMENTS

This work was supported by grants from the National Institutes of Health R01-CA164138 (SVT) and Cancer Center Support Grant P30-CA42014 as well as support from the Huntsman Cancer Foundation. BAT was supported by an NHMRC CJ Martin Early Career Fellowship (ID1091211). We acknowledge and appreciate the research participants for their continued support of our studies. We thank Wade Samowitz, MD for his consultations on tissue pathology. Research reported in this publication utilized the High-Throughput Genomics, Bioinformatic Analysis, and Genetic Counseling shared resources at Huntsman Cancer Institute and the DNA Sequencing Core Facility at University of Utah. The content is solely the responsibility of the authors and does not necessarily represent the official views of the NIH.

## DATA AVAILABILITY STATEMENT

Data that support the findings of this study are available from SVT upon request.

