## Supplemental for "A novel ribosomal protein S20 variant in a family with unexplained colorectal cancer and polyposis"

**Index of Supplemental Methods, Tables and Figures**

Supplemental Methods

Supplemental Table S1

Supplemental Table S2

Supplemental Table S3

Supplemental Figure S1

Supplemental Figure S2

**Supplemental Methods**

Subject Inclusion Criteria

CRC cases were selected for having a CRC diagnosis and at least one first-degree relative having a CRC diagnosis (n=746). Subjects known to have pathogenic variants in established CRC genes by CLIA or research testing were excluded from the study. Subjects suspected of having known hereditary cancer syndromes solely because of their clinical presentation were not excluded. Cancer-free controls (n=1,525) were selected for having: 1) no personal history of cancer, absence of any cancer in first-degree relatives, and absence of CRC in any second-degree relative (n=1353), or 2) no personal history of cancer and an absence of CRC or breast cancer in first-degree relatives (n=172).

Case-Control Sequencing and Gene Prioritization

DNA samples came either from blood (n=2094) or formalin-fixed paraffin-embedded (FFPE) tissue (n=177). Samples were sheared using a Covaris E-series (Covaris, Woburn, MA) with the following settings: duty cycle 10%, intensity 5, cycles/burst 200, time 180s (for blood-derived DNA) or 360s (for FFPE-derived DNA). For DNA extracted from FFPE blocks, 2µl of Uracil-DNA Glycosylase (NEB #M0280S) were added to samples following adapter ligation and incubated for 37°C for 15 min, prior to ligation clean-up in order to reduce C>T artefacts^1^. We designed a custom 196-gene panel (Roche SeqCap EZ Choice Library #06266339001), including 59 established or candidate cancer susceptibility genes identified through literature review and 137 additional genes selected by in-house gene prioritization analyses. Individual libraries were combined into pools of 12 prior to hybridization, and then super-pooled into 72 samples per sequencing lane.

Variant discovery using the Broad Institutes’ Genome Analysis Tool Kit (GATK) Best Practices germline workflows^2^ was conducted as follows: Sequences were aligned to the human reference genome (GRCh37) using Burrows-Wheeler Alignment tool (BWA)^3^; BAM file analyses and processing were conducted using Picard (http://broadinstitute.github.io/picard) and GATK; and joint variant calling was conducted using GATK’s HaplotypeCaller.

VCFtools was used to conduct quality control to exclude bad quality or duplicated/related samples^4^. Samples in the case-control cohort that had a heterozygosity score < -0.8; missing rate > 10%; and mean depth < 20 reads were excluded from further analysis. Individuals with a kinship coefficient > 0.25 were investigated further for relatedness ^5^. The gene-disease association test with PERCH was performed with the default settings and no contribution of biological relevance^6^.

Supplemental Table S1: Primer Sequences and PCR Conditions

| **Name** | **Usage** | **Sequence (5’ > 3’)** |
| --- | --- | --- |
| RPS20exon2/3For | Genotyping PCR | CAGATGCCGGGGCGTGTAG |
| RPS20exon2/3Rev | Genotyping PCR and sequencing | AGCTTGCGCCTGTTAAGCAC |
| RPS20exon1For | RT-PCR | GAGGATTTTTGGTCCGCACG |
| RPS20exon4Rev | RT-PCR and sequencing of 370bp and 444bp bands | TGCAATGGTGACTTCCACCT |
| RPS20intron3For | Sequencing of 591bp band | TAAAGTAAATCGGAGGGGC |
| RPS20intron2For | LOH analysis PCR and sequencing | ACTGAGAGGTCTTTTATTTCCTGT |
| RPS20intron3RevB | LOH analysis PCR and sequencing | GCCGAACTCCTTAAAGAACCTGA |
| **PCR conditions for genotyping** | | |
| 95^o^C for 10 minutes; 35 cycles of 95^o^C for 30 seconds, 59^o^C for 30 seconds, and 72^o^C for 60 seconds; 72^o^C for 7 minutes; 12^o^C hold | | |
| **PCR conditions for RT-PCR** | | |
| 95^o^C for 10 minutes; 35 cycles of 95^o^C for 30 seconds, 60^o^C for 30 seconds, and 72^o^C for 60 seconds; 72^o^C for 7 minutes; 12^o^C hold | | |
| **PCR conditions for LOH analysis** | | |
| 95^o^C for 10 minutes; 35 cycles of 95^o^C for 30 seconds, 58^o^C for 30 seconds, and 72^o^C for 60 seconds; 72^o^C for 7 minutes; 12^o^C hold | | |

All primers were used at a concentration of 0.2 µM, AmpliTaq Gold® 360 Master Mix (Applied Biosystems #4398876) was used for PCR amplification reactions, and gel electrophoresis was conducted using 2.5% agarose gel containing 0.5µg/mL ethidium bromide. PCR (polymerase chain reaction); RT-PCR (reverse transcription-polymerase chain reaction); LOH (loss-of-heterozygosity)

Supplemental Table S2: List of Genes Included in the Custom CRC Gene Panel

| **Gene** | **Previously Reported**  **Cancer Susceptibility Gene** |
| --- | --- |
| ABCA10 |  |
| ACACB |  |
| ADAMTS17 |  |
| ADGRV1 |  |
| AFF4 |  |
| AIRE |  |
| AKT2 |  |
| ANK3 |  |
| APC | Yes^7,8^ |
| ARSI |  |
| ASH1L |  |
| ASUN |  |
| ATAD2B |  |
| ATG13 |  |
| ATM | Yes^9^ |
| ATP9B |  |
| ATXN1 |  |
| AXIN2 | Yes^10^ |
| BAG3 |  |
| BARD1 | Yes^11,12^ |
| BAZ1B |  |
| BAZ2B |  |
| BLM | Yes^13,14^ |
| BMPR1A | Yes^15^ |
| BPTF |  |
| BRCA1 | Yes^16^ |
| BRCA2 | Yes^17^ |
| BRIP1 | Yes^18^ |
| BRMS1 |  |
| BUB1 | Yes^19^ |
| BUB1B | Yes^20^ |
| BUB3 | Yes^19^ |
| C9orf3 |  |
| CALCA |  |
| CD276 |  |
| CDH1 | Yes^21^ |
| CDH5 |  |
| CDKN2A | Yes^22^ |
| CENPE | Yes^23^ |
| CENPN |  |
| CEP350 |  |
| CHEK2 | Yes^24^ |
| CHSY1 |  |
| CIRBP |  |
| CNGB1 |  |
| COL1A2 |  |
| COL5A1 |  |
| COMMD8 |  |
| CRYGA |  |
| CSF1 |  |
| CYSLTR2 |  |
| DCT |  |
| DICER1 | Yes^25^ |
| DIP2B |  |
| DLEC1 |  |
| DNAJC16 |  |
| DPH2 |  |
| ECI1 |  |
| EIF1 |  |
| EIF4B |  |
| ELOVL4 |  |
| ENG | Yes^26^ |
| EPB41L4A |  |
| EPC1 |  |
| EPCAM | Yes^27,28^ |
| EXO1 | Yes^29^ |
| FAHD1 |  |
| FAM114A2 |  |
| FAN1 | Yes^30^ |
| FANCM | Yes^31^ |
| FBN3 |  |
| FEN1 |  |
| FLCN | Yes^32^ |
| FLYWCH1 |  |
| FN1 |  |
| FOCAD | Yes^33^ |
| FRAS1 |  |
| FXYD6 |  |
| GALNT12 | Yes^34^ |
| GCN1L1 |  |
| GIMAP2 |  |
| GPR132 |  |
| GPX6 |  |
| GREM1 | Yes^35^ |
| HEBP2 |  |
| IFNA13 |  |
| ITPR3 |  |
| JMJD7 |  |
| KCNQ3 |  |
| KIF20B |  |
| KIF23 | Yes^23^ |
| KMT2C |  |
| KRI1 |  |
| LAMB4 | Yes^31^ |
| LAMC3 | Yes^31^ |
| LMBRD1 |  |
| LOXHD1 |  |
| LRIG1 |  |
| LYG2 |  |
| MAST2 |  |
| MDN1 |  |
| MFAP3 |  |
| MFRP |  |
| MLH1 | Yes^36^ |
| MLH3 | Yes^37^ |
| MORC3 |  |
| MPND |  |
| MRE11A | Yes^38^ |
| MSH2 | Yes^39,40^ |
| MSH3 | Yes^41^ |
| MSH6 | Yes^42^ |
| MTCL1 |  |
| MUTYH | Yes^43^ |
| MYH7B |  |
| MYO19 |  |
| N4BP3 |  |
| N6AMT1 |  |
| NBN | Yes^44^ |
| NCAPD2 |  |
| NELFA |  |
| NF1 | Yes^45,46^ |
| NTHL1 | Yes^47^ |
| NUP214 |  |
| OAF |  |
| PALB2 | Yes^48^ |
| PCDHGB5 |  |
| PIGS |  |
| PKHD1 |  |
| PLA2G7 |  |
| PLCD4 |  |
| PMS1 | Yes^49^ |
| PMS2 | Yes^49^ |
| POLD1 | Yes^50^ |
| POLE | Yes^50^ |
| POLR3A |  |
| PPARGC1A |  |
| PPIC |  |
| PTCHD3 | Yes^31^ |
| PTEN | Yes^51^ |
| RAD50 | Yes^44^ |
| RAD51C | Yes^52^ |
| RAI1 |  |
| RANBP2 |  |
| RB1 | Yes^53^ |
| RGPD3 |  |
| RGS21 |  |
| RINT1 | Yes^54^ |
| RNF151 |  |
| RNF213 |  |
| RPL22 |  |
| RPS20 | Yes^55^ |
| RYR3 |  |
| SCG5 | Yes^35^ |
| SDC3 |  |
| SEMA4A | Yes^56^ |
| SLU7 |  |
| SMAD4 | Yes^57^ |
| SPRY4 |  |
| STIM1 |  |
| STK11 | Yes^58,59^ |
| SYNE1 |  |
| SYNE2 |  |
| TAAR5 |  |
| TAF1C |  |
| TAF5 |  |
| TDRD5 |  |
| TGFB1 |  |
| TGFBR1 |  |
| TMEM204 |  |
| TNXB |  |
| TP53 | Yes^60^ |
| TPSD1 |  |
| TRAPPC8 |  |
| TREX2 | Yes^31^ |
| TRIO |  |
| TSG101 |  |
| TSHZ3 |  |
| UCMA |  |
| USHBP1 |  |
| USP6 |  |
| VPS13A |  |
| VPS13B |  |
| VPS41 |  |
| VWA3B |  |
| VWA8 |  |
| WBP4 |  |
| WWP2 |  |
| XRCC2 | Yes^61^ |
| XRCC3 | Yes^62,63^ |
| XYLT1 |  |
| YLPM1 |  |
| ZDBF2 |  |
| ZFYVE26 |  |
| ZMIZ1 |  |
| ZNF407 |  |
| ZNF98 |  |

Supplemental Table 3: RPS20 Loss-of-Function Variant Population Frequency

| RPS20 Variants | Observed Our Study 746 cases | Observed  Our Study 1,525 controls | Observed  gnomAD v2.1.1 134,187 non-cancer^64^ |
| --- | --- | --- | --- |
| ENST00000009589:c.177+1G>A | 1 | 0 | 0 |
| ENST00000009589 Loss-of-Function Variants | 1 | 0 | 3 |
| All Transcripts Loss-of-Function Variants | 1 | 0 | 12 |

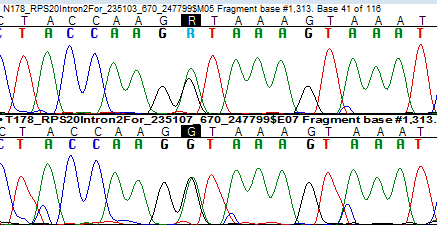
 **A**	N

T

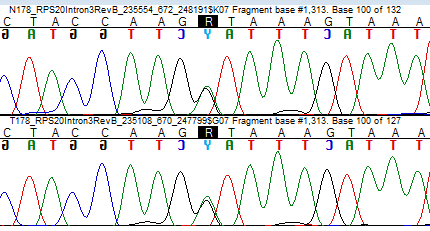
**B**	N

T

Figure S1 Sanger sequencing chromatographs of germline (N) and CRC tumor (T) DNA from *RPS20* NM_001023.3: c.177+1G>A carrier II-3. A. forward sequence B. reverse sequence

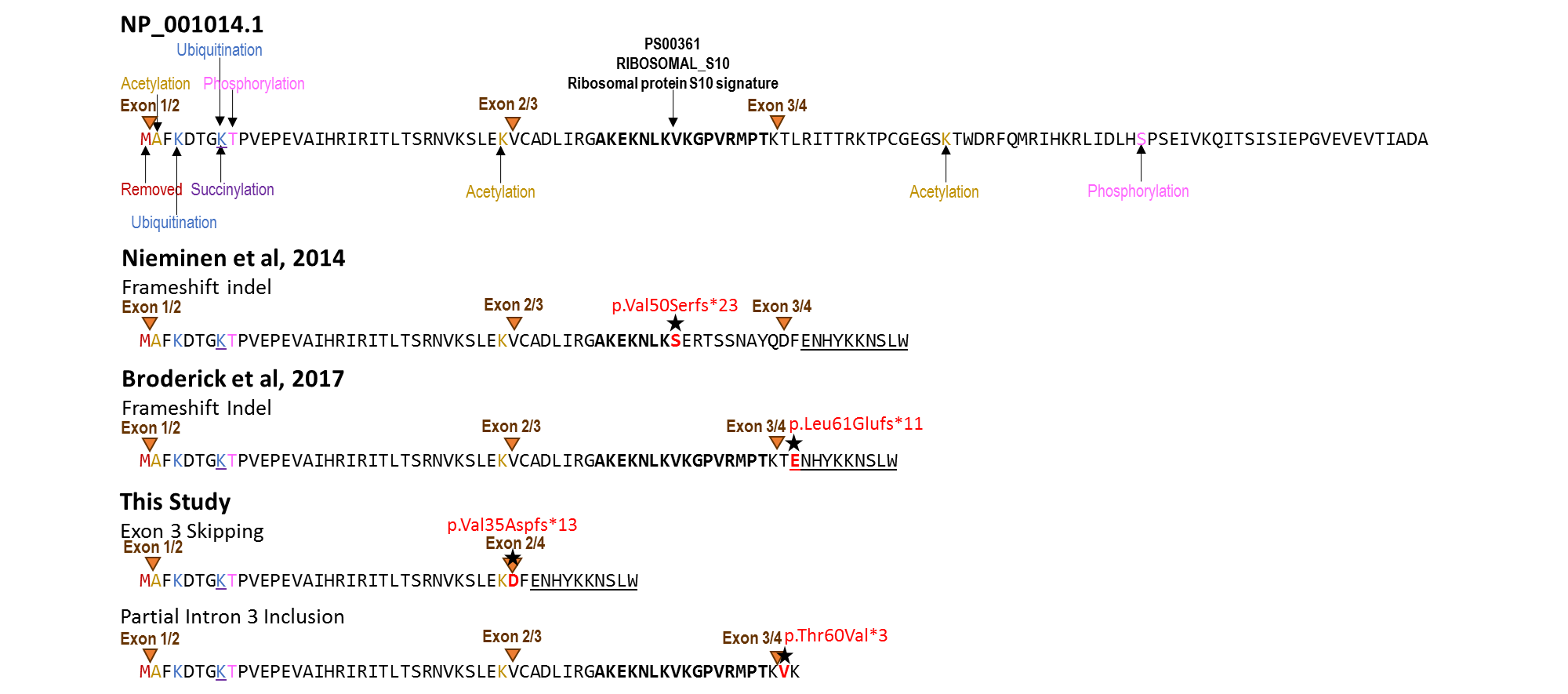

Figure S2 Predicted protein sequences of the RPS20 variants linked to colorectal cancer. Orange triangles indicate exon/exon boundaries. Stars indicate variant positions. Known and predicted protein modification sites in wild-type RPS20 NP_001014.1 indicated as follows: red text = removed, yellow text = acetylation, blue text = ubiquitination, pink text = phosphorylation, purple underline = succinylation^65^. Ribosomal protein S10 consensus pattern indicated in **BOLD**^66^.

65.	UniProt C. UniProt: a worldwide hub of protein knowledge. *Nucleic Acids Res.* 2019;47(D1):D506-D515.

66.	Sigrist CJ, de Castro E, Cerutti L, et al. New and continuing developments at PROSITE. *Nucleic Acids Res.* 2013;41(Database issue):D344-347.
